# Cytotoxicity of Activator Expression in CRISPR-based Transcriptional Activation Systems

**DOI:** 10.1101/2024.09.23.614524

**Authors:** Aakaanksha Maddineni, Ziyan Liang, Shreya Jambardi, Subrata Roy, Josh Tycko, Ajinkya Patil, Mark Manzano, Eva Gottwein

## Abstract

CRISPR-based transcriptional activation (CRISPRa) has extensive research and clinical potential. Here, we show that commonly used CRISPRa systems can exhibit pronounced cytotoxicity. We demonstrate the toxicity of published and new CRISPRa vectors expressing the activation domains (ADs) of the transcription factors p65 and HSF1, components of the synergistic activation mediator (SAM) CRISPRa system. Based on our findings for the SAM system, we extended our studies to additional ADs and the p300 acetyltransferase core domain. We show that the expression of potent transcriptional activators in lentiviral producer cells leads to low lentiviral titers, while their expression in the transduced target cells leads to cell death. Using inducible lentiviral vectors, we could not identify an activator expression window for effective SAM-based CRISPRa without measurable toxicity. The toxicity of current SAM-based CRISPRa systems hinders their wide adoption in biomedical research and introduces selection bottlenecks that may confound genetic screens. Our results suggest that the further development of CRISPRa technology should consider both the efficiency of gene activation and activator toxicity.

## Introduction

The development of programmable Clustered Regularly Interspaced Short Palindromic Repeats (CRISPR)/CRISPR-associated protein (Cas)-based transcriptional activation (CRISPRa) tools is of high interest for research and clinical applications (*1*). In these approaches, transcriptional activators are recruited to specific sites in the genome, by fusion to endonuclease-inactivated Cas proteins (“dCas”, most commonly dCas9 (*2*)) or through aptamers in the associated single guide RNA (sgRNA) (*3-7*). The appeal of CRISPRa over traditional cDNA expression approaches lies in its simplicity of sgRNA design, scalability, ability to multiplex, and ability to overexpress relevant gene isoforms and large transcripts from their endogenous loci. CRISPRa can furthermore target non-coding genes and regulatory loci for transcriptional activation. CRISPRa is most commonly achieved by the fusion of dCas9 with the transactivation domains (ADs) of transcription factors (TFs), which promote transcription by, in turn, recruiting transcriptional and epigenetic machinery, including general transcription factors, the Mediator complex, and chromatin-modifying enzymes. The first generation of CRISPRa vectors used several copies of an 11 amino acid peptide representing the minimal activation domain (AD) of the herpes simplex virus type 1 TF virion protein 16 (VP16) (*3, 5, 7*). More potent second-generation activators relied on recruiting additional ADs (*4, 6, 8-10*), either by fusion to dCas9 or through bacteriophage RNA-aptamers engineered into the scaffold portion of the sgRNA, or both. Among the most potent and commonly used CRISPRa approaches is the synergistic activation mediator (SAM) system (*4, 10*) (Fig. 1A). In the SAM system, dCas9 is fused to four copies of the VP16 minimal AD (VP64) and loaded with an aptamer-modified sgRNA which in turn recruits the MS2 or PP7 bacteriophage coat protein (MCP/PCP)-fused ADs of p65 or HSF1. We will abbreviate MCP or PCP-fused p65^AD^-HSF1^AD^ synthetic transcriptional activators as MPH and PPH here. The original SAM system consists of 3 lentiviral vectors (LVs) (*4*), expressing dCas9-VP64, MPH, and the aptamer-modified sgRNA, respectively. A subsequent version of SAM aimed to improve the titer of the original MPH-encoding LV and reduce the number of required LVs to two vectors (*11*) by combining a PPH activator protein and an aptamer-modified sgRNA in a single vector (pXPR_502, Fig. 1B). In a conceptually different approach (*12*), with potentially distinct target preferences (*13*), CRISPRa is achieved through dCas9- or sgRNA-mediated recruitment of histone or DNA-modifying enzymatic domains (*12*), such as the catalytic histone acetyltransferase (HAT) core domain of the human E1A-associated protein p300 (*14*).

**Fig. 1.**
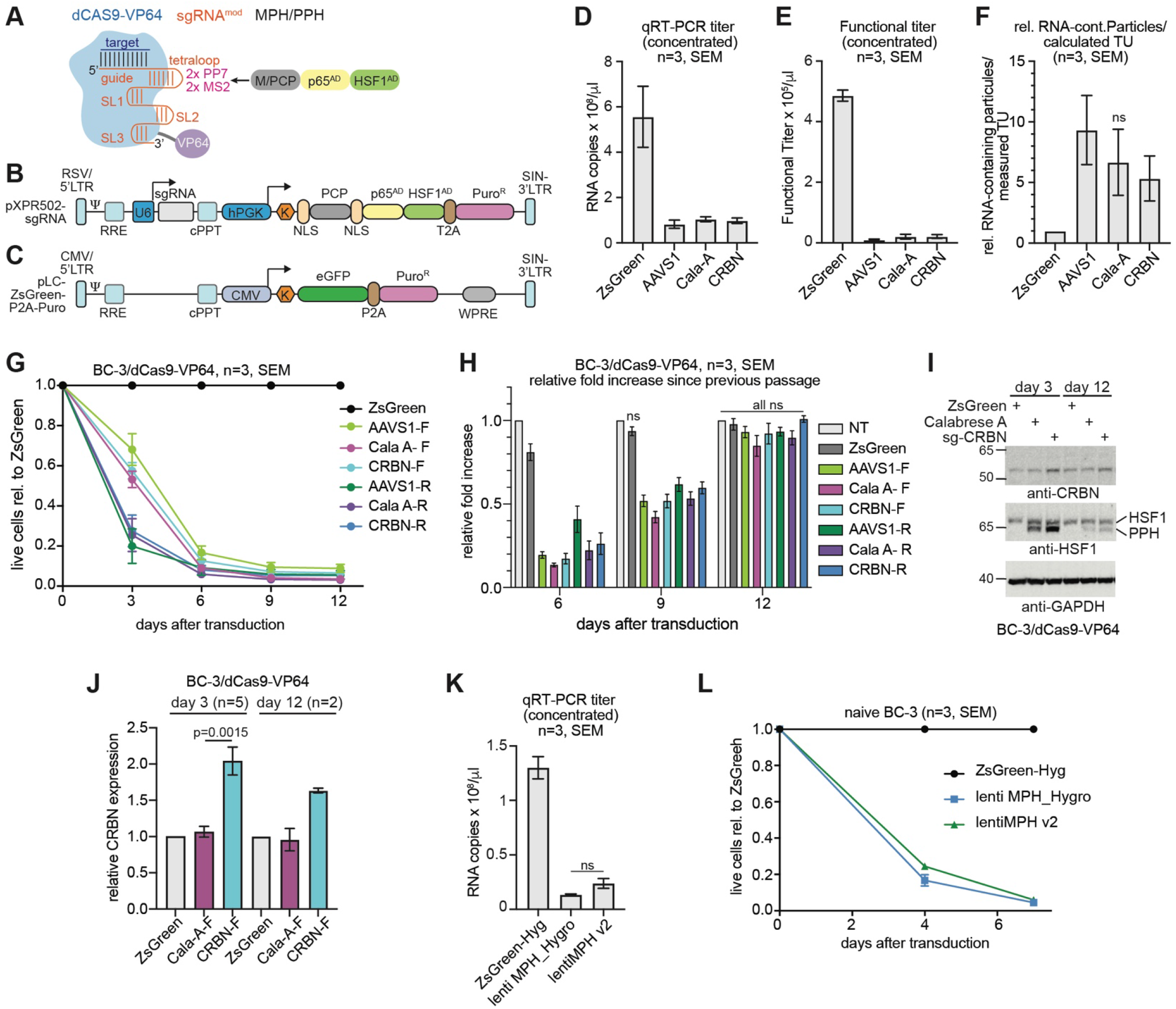
Published p65^AD^-HSF1^AD^-expressing LVs have low titers and result in lower-than-expected outgrowth after transduction. **(A)** Schematic of the SAM CRISPRa system. sgRNA features are in orange. In pXPR_502, the sgRNA tetraloop is modified by two PP7 and two MS2 aptamers. **(B)** Schematic of the pXPR_502 lentiviral vector, showing the long-terminal repeats (LTRs), the Rev responsive element (RRE), the packaging signal (Ψ), the U6 promoter, the sgRNA cassette, the PGK promoter driving a nuclear PPH-T2A-Puro^R^ fusion protein, the central polypurine tract (cPPT). SIN: self-inactivating. K: Kozak sequence. Not drawn to scale. **(C)** Schematic of pLC-ZsGreen-P2A-Puro, not drawn to scale, aligning similar components to panel B. **(D)** RNA titers of concentrated lentiviral stocks generated from pLC-ZsGreen-P2A-Puro, or pXPR_502 expressing sgAAVS1 (AAVS1), Calabrese Set A, or sgCRBN-a1 (CRBN). Data are from 3 independent stocks per LV. All titers were significantly different from that of the ZsGreen control, unpaired t test, p<0.05. Error bars represent SEM. **(E)** Functional titers in unmodified BC-3 of the same LV stocks as in panel C, read on day 2 after transduction, after one day of puromycin selection. All titers were significantly different from that of the ZsGreen control, unpaired t test, p<0.05. Error bars represent SEM **(F)** Relative ratio of RNA-continaining LV particles to measured transducing units (TUs), calculated from values in C and D, assuming 2 LV-gRNAs per LV particle. Values were significantly higher than for the ZsGreen control, unpaired one-sided t test, p<0.05, except for Cala-A, as indicated by ns. The Cala-A value was significantly higher (p<0.05) using a ratio-paired t-test. Error bars represent SEM. **(G)** Growth curve analyses of pXPR_502-transduced BC-3/dCas9-VP64. Shown are cumulative cell counts relative to ZsGreen-P2A-Puro (ZsGreen)-control transduced cells. Samples were transduced at a calculated MOI 0.3, based on either functional (F) or RNA (R) titers, relative to the titer of pLC-ZsGreen-P2A-Puro. 3 independent repeats, using the 3 LV preps from panels D-F. For results from naïve BC-3, see Fig. S1A. All values differed significantly from the ZsGreen control, unpaired t test, p<0.05, n=3 independent repeats. Error bars represent SEM. **(H)** Fold increase over the previous passage on days 6, 9, and 12, from the same experiments shown in panel G. Values were normalized to the untransduced and unselected control samples (NT) in this analysis. For results from naïve BC-3, see Fig. S1B. Values were significantly different from NT at each time point unless specified by ns, unpaired t test, p<0.05. Error bars represent SEM. **(I)** Western Blot analyses of CRBN, PPH, and GAPDH expression in representative lysates taken on day 3 or day 12 after transduction from a subset of the samples shown in Fig. 1G-H. Endogenous HSF1 is marked “HSF1”. For quantification over 5 (day 3) or 2 (day 12) independent repeats, see Figs. 1J and S1D. For results from BC-3, see Figs. S1A-C. **(J)** Quantification of results shown in Fig. 1I. Data from 5 (day 3) or two (day 12) independent repeats. Unpaired t test. Error bars represent SEM. **(K)** As in Fig. 1D, but using a hygromycin resistant control vector (pLC-ZsGreen-P2A-Hyg) and lenti MS2-P65-HSF1_Hygro (*4*) or lentiMPH v2 (*20*). Data are from three independent LV preparations. Values were significantly different from the ZsGreen control, unpaired t test, p<0.05. Differences between the two MPH vectors were not significant (ns). Error bars represent SEM. **(L)** As in Fig. 1G but using LVs from Fig. 1K and ending the growth curve on day 6. One repeat with each LV preparation, n=3 repeats overall. Results on day 4 and 6 differed significantly from the control, unpaired t test, p<0.05. Differences between the two MPH vectors were not significant at either time point. Error bars represent SEM.

We attempted to establish the SAM system for our work in primary effusion lymphoma (PEL) B cell lines, a cell model we have extensively used in CRISPR/Cas9 screens (*15-17*). During these experiments, we encountered difficulties with LVs encoding MPH or PPH fusion proteins, including apparently low lentiviral titers and an inability to obtain transduced cell pools at expected efficiencies in several different PEL cell lines. Here, we systematically investigated the technical barriers to adapting CRISPRa for our work. Our results suggest that both published and novel vectors expressing activators used for CRISPRa exhibit pronounced cytotoxicity, leading to low lentiviral titers and target cell death. This toxicity could confound studies using CRISPRa and should be considered in the further development of this technology.

## Results

### SAM activation domain vectors are toxic under conditions used for screening

To investigate the reasons for our inability to implement robust protocols for SAM-based CRISPRa in PEL cell lines, we initially focused on a set of pXPR_502 vectors (*11*) (Fig. 1B), expressing the PPH activator fusion protein and either the genome-scale Calabrese sgRNA library set A (*11*) (Cala-A) or individual sgRNAs targeting the safe harbor locus adeno-associated virus integration site 1 (AAVS1) or the promoter of the non-essential gene cereblon (CRBN), which we have stably overexpressed in PEL cells using a lentiviral cDNA vector in an unrelated study (*18*). As a control, we used an LV expressing a ZsGreen-P2A-Puro^R^ cassette (*17*) (Fig. 1C). All transfer vectors used in this study were well below the size limit for efficient packaging of HIV-based LVs (see below). We titered LV stocks by qRT-PCR and functionally in the PEL cell line BC-3 (*19*). For functional titration, we counted the percentage of cells that survived puromycin selection for all vectors and additionally used flow cytometry for the ZsGreen vector (see Methods). pXPR_502 vector preparations had lower qRT-PCR-based titers than the LV expressing ZsGreen (Fig. 1D), despite using an optimized amount of transfer vector during pXPR_502 packaging (see Methods and compare relative titers in Fig. 1D to those in Fig. 2D, where we used a standard packaging protocol). The discrepancies in the calculated functional titers were even greater than for the genomic RNA (LV-gRNA) content (Fig. 1E-F), which could result from a loss of transduced cells due to MPH toxicity soon after transduction, thereby confounding results from antibiotic selection.

**Fig. 2.**
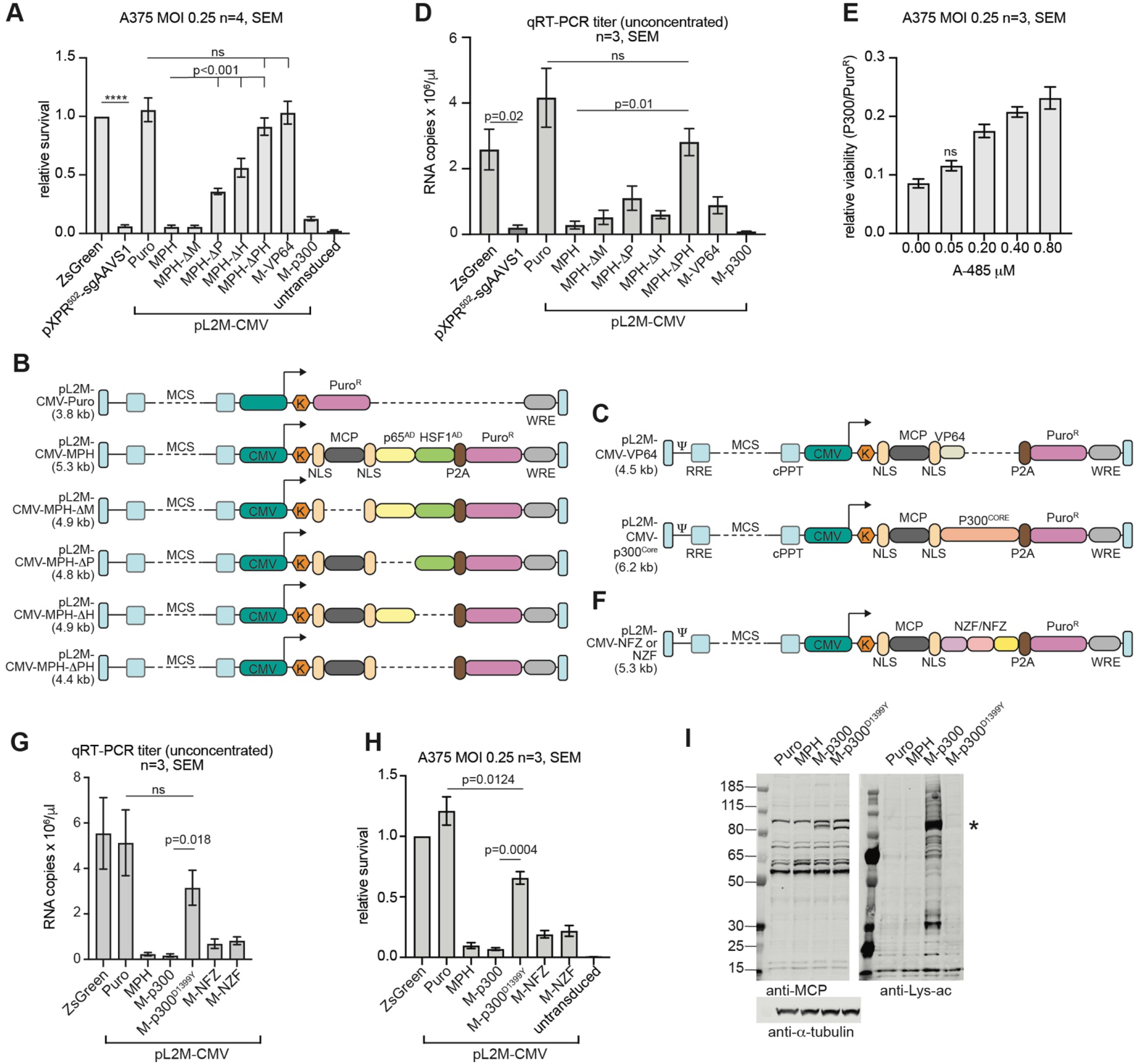
Toxicity results from the expression of strong ADs or the p300^Core^ domain. **(A)** Relative survival of A375 cells following puromycin selection after transduction with pLC-ZsGreen-P2A-Puro at MOI 0.25 based on functional titration and other LVs based on LV-gRNA content relative to pLC-ZsGreen-P2A-Puro. Cells were split (1:2) twenty-four hours after transduction and selected with puromycin. On day 3 after transduction, survival was measured using CellTiter-Glo 2.0 and normalized to the ZsGreen control. Four independent repeats using at least three independent LV stocks per vector. Results from pL2M-CMV vectors differed significantly from the matched CMV-Puro control unless indicated by ns, unpaired t-test, p<0.01. All samples had significantly higher viability than untransduced and selected controls (selected). Differences between MPH and the ΔP, ΔH, and ΔPH deletion mutants were significant (p<0.001), as were differences between ΔP or ΔH and ΔPH (p<0.02). **** denotes p<0.0001. Error bars represent SEM. **(B-C)** Schematics of the LV vectors used in Fig. 2A and D, not drawn to scale. Components are labeled as in Fig. 1B. Numbers at left indicate the distance from the transcription start site to the polyA signal in kb, showing that each LV-gRNA is several kb below the ∼9.2kb HIV genome. Distances were rounded up to the next decimal. Dashed lines represent sequences that are absent compared to the other vectors. **(D)**RNA titers of unconcentrated lentiviral stocks used in Fig. 2A, three independent virus stocks per vector. In this experiment, all vectors were packaged using 45% transfer vector (see Methods, resulting in a significantly greater discrepancy between pXPR502-sgAAVS1 and the ZsGreen control titers than in Fig. 1D, where 15% transfer vector was used for pXPR502, unpaired t test, p=0.04). Titers of all pL2M-CMV vectors differed significantly from the CMV-Puro control, except for MPH-ΔPH, unpaired t test, p<0.05, n=3. Other p values are indicated in the figure. Error bars represent SEM. **(E)**The relative survival of pL2M-CMV-MCP-p300^Core^-transduced cells compared to that of pL2M-CMV-Puro-transduced cells increased upon treatment with the HAT inhibitor A-485. Cells were transduced and assayed as in panel A. Survival compared to the DMSO-treated control was significantly increased (unpaired t test, p<0.003, n=3), except for the lowest A-485 concentration, as indicated by ns. Error bars represent SEM. **(F)**Schematic of the LV used to express the MCP-NFZ/NZF fusion proteins in panels G-H. **(G)**RNA titers for additional LVs, as in panel Fig. 2D. Titers of all pL2M-CMV vectors were significantly different from the Puro control, except for M-p300^D1399Y^, unpaired t test, p<0.05, n=3. Other p values are indicated in the figure, ns not significant. Error bars represent SEM. **(H)**Relative survival of A375 cells after transduction at MOI 0.25, as in panel A. Survival of all pL2M-CMV transduced samples was significantly different from the Puro control, unpaired t test, p<0.05, n=3. All samples had significantly higher viability than untransduced and selected controls (selected). **(I)**Western Blot analysis of MCP fusion protein expression and lysine acetylation two days after transduction of A375 at MOI 0.25, without selection, but otherwise as in Fig. 2H. We note that off-target acetylation is detected against the background of a majority of untransduced cells in this experiment, due to low MOI transduction. The band marked by an asterisk likely represents MCP-p300 autoacetylation.

To further quantify this effect, we transduced BC-3 expressing dCas9-VP64 or parental BC-3 at a multiplicity of infection (MOI) of ∼0.3, either based on the calculated functional titer of each vector (“F”) or based on LV-gRNA copy numbers relative to the ZsGreen-expressing control vector (“R”). Similar MOIs are typically used in CRISPR screens to ensure delivery of one sgRNA per transduced cell. We selected the resulting cell pools using puromycin and performed growth curve analyses. For both cell lines, a close to expected fraction of the ZsGreen control vector-transduced cells survived puromycin selection (∼39% compared to untransduced and unselected control cells on day 3 after transduction, ∼6.5% range) and proliferated similarly to untransduced cells once puromycin selection was complete (Fig. 1G-H, S1A-B). In contrast, dramatically fewer pXPR_502-transduced cells survived over time for either titration approach, likely indicating ongoing transgene toxicity (Fig. 1G-H, Fig. S1A-B). This toxicity was independent of the specific sgRNA insert and the presence of dCas9-VP64 and associated gene activation, which we readily observed for sgCRBN (Fig. 1I-J). After continued passage, pXPR_502-transduced cell pools that proliferated normally were obtained by about day 9 after transduction, demonstrating that it is possible to obtain cells that have overcome PPH toxicity (Fig. 1H, S1B). Western blot analyses show that these passaged cell pools had ∼5-fold reduced expression levels of PPH (Figs. 1I, Fig. S1C-D) and, therefore, either contain cells with low initial PPH expression or those that have undergone changes resulting in lower PPH expression. These cell pools maintained a reduced level of CRISPRa-based gene activation (Fig. 1I-J). Low titer and severe toxicity in BC-3 cells were also evident with commonly used MPH-encoding vectors for the 3-LV SAM system (*4, 20*), showing that our findings are not exclusive to pXPR_502 (Fig. 1K-L). In these experiments, differences between the original (lenti MS2-P65-HSF1_Hygro (*4*)) and an updated (lentiMPH v2 (*20*)) MPH vector did not reach statistical significance over three independent virus preparations and transductions.

### Several MCP-Fused CRISPRa Activators are Toxic Across Contexts

We next tested whether these observations were unique to the PEL model by repeating the experiment for pXPR_502-sgAAVS1 in the melanoma cell line A375, which was used in several published CRISPRa screens (*4, 11*). Following transductions at ∼MOI 0.25, based on LV-gRNA content and the functional titer of the ZsGreen-expressing positive control vector, pXPR_502-toxicity was also pronounced in A375, suggesting that toxicity is not unique to the PEL model (bars 1 and 2 in Fig. 2A). As for BC-3, we were able to grow out pXPR502-transduced A375 cell lines after a severe bottleneck (not shown).

To map potentially cytotoxic components of the M/PCP-p65^AD^-HSF1^AD^ fusion proteins, we constructed a lentiviral vector expressing only MCP-p65^AD^-HSF1^AD^ (MPH), versions lacking MCP (ΔM), p65^AD^ (ΔP), HSF1^AD^ (ΔH), or p65^AD^-HSF1^AD^ (ΔPH), and a matched control vector expressing only the puromycin resistance gene (CMV-Puro, for vector schematics see Fig. 2B and for expression controls see Fig. S2). We finally constructed matched vectors expressing fusions of MCP with VP64 or the HAT core domain of p300 (p300^Core^) (Fig. 2C), reasoning that these vectors could eventually be used together with dCas9-p300^Core^ or dCas9-VP64, respectively, as proposed previously (*21*). All fusion proteins were targeted to the nucleus using dual nuclear localization signals (NLS) flanking MCP, like in pXPR_502. After normalizing for LV-gRNA copies relative to the ZsGreen control, the CMV-Puro LV achieved the expected numbers of transduced cells, validating our titration strategy (Fig. 2A). In contrast, few cells survived transduction with the MPH-encoding LV. Deletion of each AD partially rescued vector toxicity and deletion of both ADs eliminated toxicity.

Interestingly, p65^AD^-HSF1^AD^ deletion also rescued LV titers, suggesting that the low titers of MPH vectors likely result from AD toxicity in the producer cells (Fig. 2D). MCP-VP64-transduced cells did not experience toxicity (Fig. 2A) consistent with our ability to establish dCas9-VP64-expressing PEL cell lines. The lack of VP64 toxicity in A375 could be due to its weaker activator activity and lower levels of MCP-VP64 expression (Fig. S2B-C). In contrast, MCP-p300 expression was toxic in A375 (Fig. 2A). MCP-p300^Core^-transduction at single copy resulted in a readily detectable increase in lysine acetylation, including autoacetylation of MCP-p300^Core^ (Fig. S3 and below), suggesting an sgRNA-independent off-target activity of this fusion protein. MCP-p300^Core^-induced lysine acetylation was partially reversed by treatment with increasing concentrations of A-485 (Fig. S3), an inhibitor of the catalytic activity of p300/CBP (*22*). A-485 treatment furthermore significantly, albeit partially, rescued the survival MCP-p300^Core^-transduced cells (Fig. 2E). While the toxicity of A-485 precluded testing higher concentrations, the substantial rescue of MCP-p300^Core^ toxicity by partial HAT inhibition further validates our titration approach and shows that the toxicity of this vector is at least partially due to p300 HAT activity.

Using a genetic approach, we constructed a p300^Core^ mutant with inactivated HAT activity (p300^Core^/D1399Y) (*14*). In the same experiment, we tested MCP fusions with the recently reported NFZ and NZF triple AD cassettes (*23*), containing the ADs of the TFs NCOA3 (N), FOXO3 (F), and ZNF473 (Z) in a different order (Fig. 2F). While NFZ and NZF were previously only tested following direct dCas9-mediated recruitment, we tested them as MCP fusion proteins, which could eventually improve AD potency during CRISPRa through multicopy recruitment through MS2 stem-loops, while mitigating MPH toxicity. NFZ/NZF fusion to MCP furthermore enables a direct comparison to other constructs in this figure. As expected, based on the A-485-mediated rescue, the p300^Core^/D1399Y mutation rescued LV titers and strongly reduced p300^Core^ toxicity (Fig. 2G-H). p300^Core^/D1399Y-mutation additionally abolished off-target acetylation and autoacetylation of the MCP-p300^Core^ (Fig. 2I). MCP-NZF and MCP-NFZ constructs had low titers and measurable toxicity in this experimental context, showing that these new activators are unlikely to overcome the toxicity of current CRISPRa systems completely. Retesting a subset of the vectors in this figure by transduction of BC-3 cells confirmed the reduced toxicity after AD deletion or p300 mutation and MCP-NFZ/NZF toxicity in a second cellular context (Fig. S4).

### Inducible MPH expression is toxic in various cell lines

In the experiments shown above, toxicity occurred already during the antibiotic selection of the transduced cells, making it difficult to distinguish low titer from transgene toxicity. Our finding that cells that survive the toxicity bottleneck have reduced activator expression (Figs. 1H-J, S1B-D) suggests that high levels of activator expression are toxic, while lower expression levels might be tolerated. To uncouple antibiotic selection from transgene toxicity and allow for tunable activator expression, we constructed a doxycycline-inducible expression vector for MPH and established cell lines based on BC-3, BC-3/dCas9-VP64, A375, the T cell line Jurkat, and 293T by LV transduction at single copy. We observed dose-dependent doxycycline-induced toxicity upon MPH induction in each cell line (Figs. 3, S5), showing that toxicity can be uncoupled from LV transduction and suggesting that the activation domains of at least p65^AD^ and HSF1^AD^ are toxic across an expanded set of cellular contexts.

**Fig. 3.**
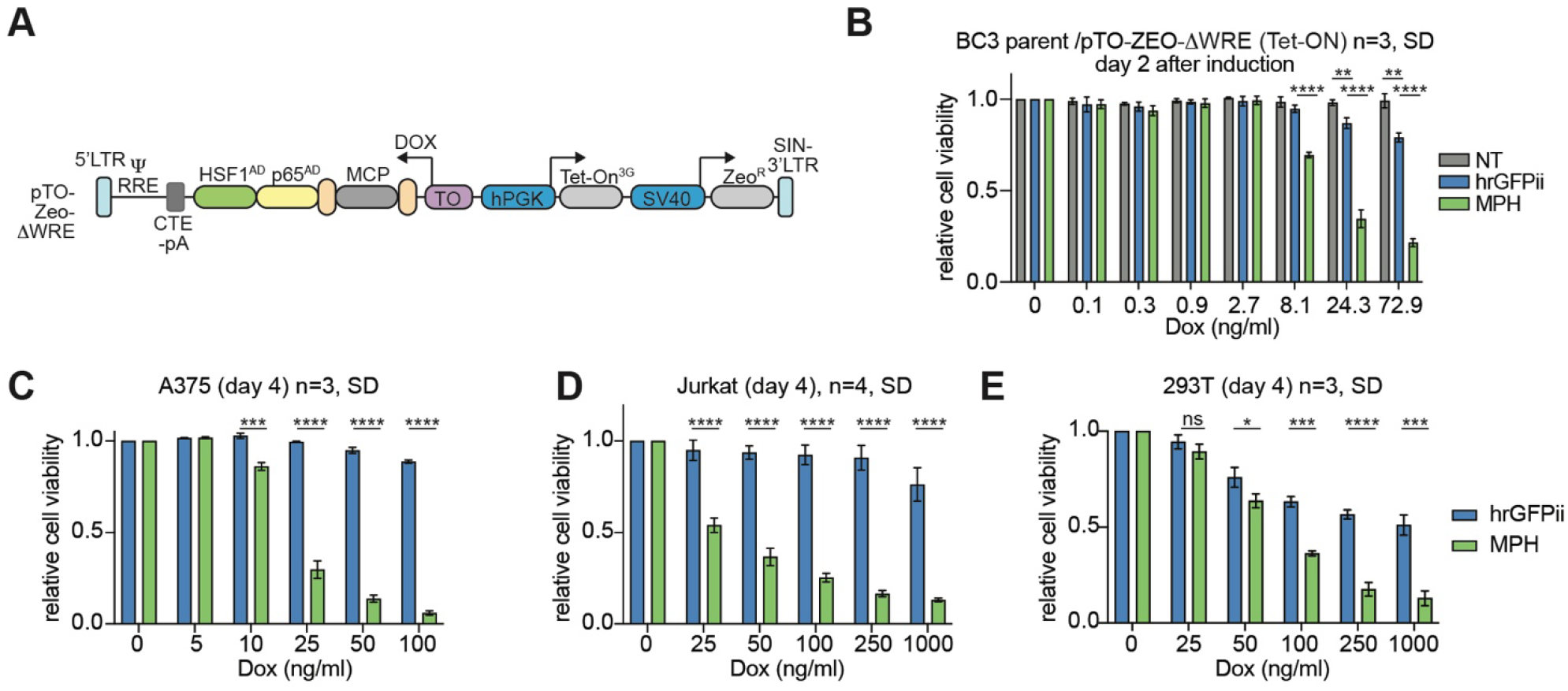
Inducible MPH expression is toxic in various cell lines. **(A)**Schematic of the Dox-inducible lentiviral vector pTO-Zeo-ΔWPRE. **(B)**Relative survival of untransduced (NT) BC-3, or BC-3 expressing Dox-inducible hrGFP2 or MPH, 50 hours into treatment with the indicated concentrations of Dox. **(C-E)** As in B, but in A375, Jurkat, and 293T, respectively. Throughout, **** denotes p<0.0001, *** p<0.001, ** p<0.01, * p<0.05, and ns “not significant”, unpaired t-tests, n=3-4 as indicated. Error bars represent SD over 3-4 biological repeats.

### The SAM system is unlikely to allow efficient CRISPRa without measurable toxicity

Leveraging the inducible MPH expression vector, we tested whether there is a window of MPH expression that allows for CRISPRa without measurable toxicity or prior adaptation to MPH expression. Based on our result that toxicity was measurable 2 days after induction with 8ng/ml Dox, we treated BC-3/dCas9-VP64/Tet-ON-MPH cells with concentrations between 2 and 8 ng/ml Dox. This resulted in a window of MPH expression with a ∼45-fold range two days into induction (Figs. 4A-B, S6A). The increased MPH expression was accompanied by increasing toxicity (Fig. 4C, S6B). Based on these data, we additionally transduced the cells with vectors expressing sgAAVS1 or sgCRBN a1 (see Fig. S6C for a schematic). This approach resulted in a modest CRISPRa-mediated overexpression of CRBN upon induction of MPH, but not upon hrGFP2, despite the expression of VP64 (Fig. 4D-E). The overexpression of CRBN reached significance at 5ng/ml DOX, a concentration that killed 40% of the culture within 6 days of induction. This result suggests that the SAM system is unlikely to allow for robust CRISPRa without any measurable toxicity.

**Fig. 4.**
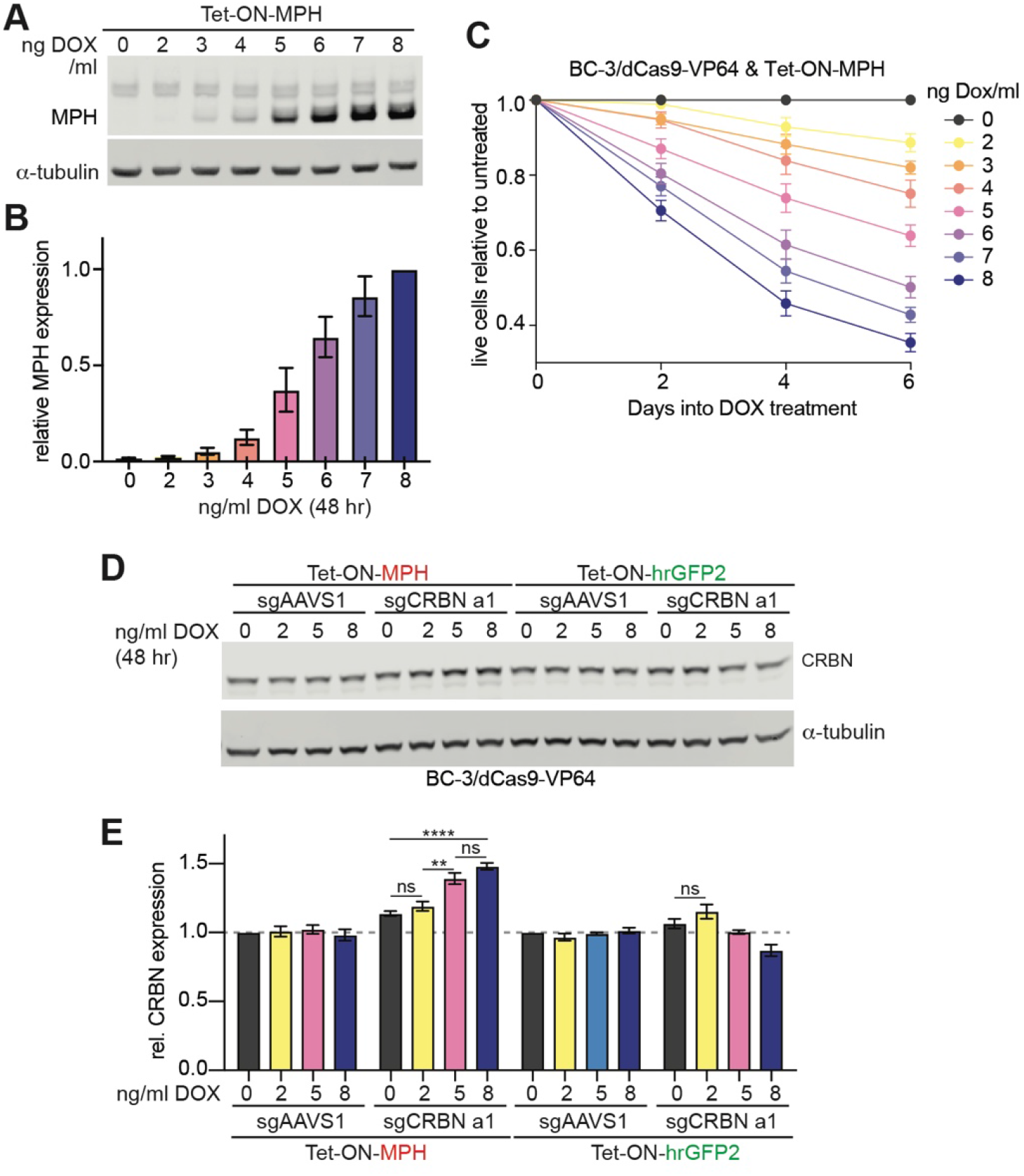
The SAM system is unlikely to allow efficient CRISPRa without measurable toxicity. **(A)**Western Blot analysis of MPH expression, using anti-HSF1, 2 days into Dox-induction. **(B)**Quantification of results from panel A over n=3, MPH expression was sequentially normalized to α-tubulin and the normalized intensity for 8ng/ml DOX. **(C)**Growth curve analysis of BC-3/Tet-ON-MPH after treatment with Dox at the indicated concentrations. Toxicity reached significance with 5ng/ml on day 2 and 2 ng/ml on days 4 and 6. Differences between 2, 5, and 8 ng/ml Dox-treated cells were significant on days 4 and 6. **(D)**Western Blot analyses of CRBN and α-tubulin expression in representative lysates taken on day 2 after induction of BC3/dCas9-VP64/Tet-ON-MPH or -hrGFP2 that were additionally transduced with sgAAVS1 or sgCRBN a1. **(E)**Quantification of results shown in 4D over 4 independent repeats. **** denotes p<0.0001, *** p<0.001, ** p<0.01, * p<0.05, and ns “not significant”, unpaired t-tests, error bars represent SEM.

## Discussion

Our data show that ectopic expression of transcriptional activator cassettes that are commonly used for CRISPRa is cytotoxic. For the SAM system, toxicity is pronounced even when published lentiviral vectors are delivered at single copy into cell types that were previously used for CRISPRa screens, such as A375. Activator domain toxicity is likely responsible for the low titers of these LVs, since AD deletions that rescued toxicity after transduction also rescued RNA-based LV titers. While MPH-transduced cells can be grown out, the observed vector toxicity represents a strong perturbation, and the bottleneck these cells experience likely affects screening results, particularly when MPH is delivered together with the sgRNA. Activator toxicity can additionally cause functional titrations to substantially underestimate LV titer (Fig. 1F), which could result in the unintentional delivery of several sgRNA per cell during screens, thereby further confounding results.

Although the literature reports the low titer of LVs used for CRISPRa (*11, 20*), there are few reports of CRISPRa off-target toxicity, perhaps because cells can eventually be grown out from the initial bottleneck. In flies, fusion of either MCP or dCas9 to the catalytic domain of the HAT CBP was reported to result in male sterility and off-target lysine acetylation (*13*), similar to Figs. 2I and S3. A recent report mentions an inability to establish cell lines expressing dCas9 fused to VP64, p65, and Epstein-Barr virus RTA ADs (dCas9-VPR) (*9, 23*).

While our experiments have not addressed the mechanisms of CRISPRa vector toxicity directly, ADs of viral or cellular TFs may compete with endogenous transcription factor complexes for cofactors that are limiting, in a process reminiscent of “cofactor squelching” for example by VP16 (*24-26*). It is also possible that AD-fusion proteins are recruited to and interfere with the function of endogenous TF complexes. Since the weak activator VP64 was well tolerated after transduction, it appears likely that activator strength correlates with toxicity, although there may be cell type-specific differences, as suggested by the imperfect correlation between titers of 293T-derived vector stocks (Fig. 2B) and toxicity in A375 (Fig. 2A).

While CRISPRa by recruitment of enzymatic domains could potentially overcome toxicity due to cofactor competition by highly expressed ADs, LVs expressing fusion proteins of MCP with the p300^Core^ domain were also toxic, at least partially due to unintended acetylation events. Rescue from toxicity upon AD deletion or p300^Core^-HAT inactivation suggests toxicity is unlikely due to competition for the nuclear import machinery since all constructs contained two NLS motifs in each fusion protein. Further studies of the mechanisms underlying CRISPRa toxicity might identify strategies to overcome the limitations of current CRISPRa systems and inform our understanding of basic concepts of transcriptional regulation, including physiological competition for limiting cofactors. Our results also underscore the importance of characterizing the recruited factors for each AD and developing additional ADs for CRISPRa (*23, 27, 28*).

We speculate that difficulties implementing CRISPRa in the broader community may be limiting to the wide adaptation of this technology. The development of this technology should therefore include assessing the toxicity of CRISPRa vectors and testing strategies to limit this toxicity. Since transduced cells that grow out after passage had strongly reduced p65^AD^-HSF1^AD^ expression, one strategy to overcome or manage CRISPRa toxicity could be to reduce the expression of AD-fusion proteins, for example by avoiding unnecessary codon optimization, using weaker or inducible promoters, or omitting sequences that boost gene expression from lentiviral vectors, such as the Woodchuck Hepatitis Virus posttranscriptional regulatory element (WPRE). Our results with inducible MPH vectors, however, suggest that it might be difficult to identify tolerated expression levels that allow for efficient CRISPRa in the absence of toxicity or selecting surviving cells, as we have done in Fig. 1.

In principle, inducible dCas9-triple activator fusions, including dCas9-VPR or dCas9-NFZ, could result in more efficient complex assembly due to the requirement for only two complex components (dCas9-AD fusion protein and sgRNA) compared to the assembly of three components in the SAM system. These dCas9 fusion proteins may also be less well-expressed and therefore less toxic than smaller MCP-AD fusions. Aptamer-mediated recruitment in contrast offers the potential for multicopy recruitment of MCP-AD fusions at lower expression level. Regardless of the approach, monitoring and controlling for AD toxicity after transduction is likely easier in CRISPRa systems where AD-expressing cell lines are established and validated first, followed by delivery of only the sgRNA during screening. Experimentally evolving the CRISPRa machinery for more efficient dCas9-AD-sgRNA-target complex assembly or reduced toxicity represents a final strategy.

Until CRISPRa systems with less pronounced toxicity are developed, best practices for performing CRISPRa experiments and screens we recommend include (i) performing titrations of constitutive AD-expressing LVs by qRT-PCR relative to a non-toxic vector, like in Figs. 1 and 2, or by including a fluorescent marker that can be analyzed before evident toxicity, (ii) establishing and validating activator-expressing cell lines before delivery of sgRNAs, while monitoring vector toxicity, and (iii) validating any screening results in unmodified cell lines using orthogonal methods, such as cDNA expression. While these approaches may help control for the toxicity of CRISPRa in a laboratory setting, it could be more difficult to control and overcome the toxicity of CRISPRa in clinical applications.

In sum, while CRISPRa remains a conceptually appealing approach, our work reveals the importance of measuring activator domain toxicity of CRISPRa systems. Our results also underscore the importance of understanding mechanisms of AD action and CRISPRa off-target toxicity, developing additional CRISPRa activators, and designing approaches that overcome toxicity.

Our study seeks to point out and characterize an important caveat of current CRISPRa technologies. Limitations of this study include that we have focused on the SAM system, NFZ/NZF, and the p300^Core^ domain and have not tested all published activators, leaving, for example, VPR and the CBP HAT domain for future investigation. We have limited our study of published vectors to the most used subset. Our experiments do not include in-depth mechanistic investigations of the observed toxicity, and we have explored only a subset of approaches one could use to overcome the toxicity of current CRISPRa approaches experimentally.

## Materials and Methods

### Experimental Design

AM, SJ, and EG specifically designed this study to quantify and characterize the unexpected performance of lentiviral vectors and resulting cell lines expressing CRISPRa components. These initially “anecdotal” effects included poor titer, unexpectedly poor or no outgrowth of transduced cell lines, and failure to recover established cell lines after storage in liquid nitrogen.

### Cell Culture

293T/17 (“293T”) (ATCC, CRL-11268) were grown in Dulbecco’s Modified Eagle’s Medium (DMEM, Corning, 10-017-CV) containing 10% Serum Plus™ II Medium Supplement (Sigma-Aldrich 14009C-500ML, Batch Number: 21C421) and 10 μg/ml gentamicin. A375 (ATCC, CRL-1619) were grown in DMEM containing 10% fetal bovine serum (FBS, Corning, 35-010-CV) and 10 μg/ml gentamicin (Gibco, 15710072). BC-3 (ATCC, CRL-2277) were grown in RPMI (Corning, 10-040-CV), containing 20% FBS and 10 μg/ml gentamicin.

### Published Constructs

The Human Calabrese CRISPR activation pooled library set A (*11*) was a gift from David Root and John Doench (Addgene #92379). pXPR_502 (*11*) was a gift from John Doench & David Root (Addgene plasmid # 96923; http://n2t.net/addgene:96923; RRID:Addgene_96923). pLX-sgRNA (*29*) was a gift from Eric Lander & David Sabatini (Addgene plasmid # 50662; http://n2t.net/addgene:50662; RRID:Addgene_50662). lentiGuide-Puro (*30*) was a gift from Feng Zhang (Addgene plasmid # 52963). pcDNA-dCas9-p300 Core (*14*) was a gift from Charles Gersbach (Addgene plasmid # 61357; http://n2t.net/addgene:61357; RRID:Addgene_61357). Lenti dCAS-VP64_Blast (*4*) was a gift from Feng Zhang (Addgene plasmid # 61425; http://n2t.net/addgene:61425; RRID:Addgene_61425). lenti MS2-P65-HSF1_Hygro (*4*) was a gift from Feng Zhang (Addgene plasmid # 61426; http://n2t.net/addgene:61426; RRID:Addgene_61426). lentiMPH v2 (*20*) was a gift from Feng Zhang (Addgene plasmid # 89308; http://n2t.net/addgene:89308; RRID:Addgene_89308). pMD2.G was a gift from Didier Trono (Addgene plasmid # 12259; http://n2t.net/addgene:12259; RRID:Addgene_12259). psPAX2 was a gift from Didier Trono (Addgene plasmid # 12260; http://n2t.net/addgene:12260; RRID:Addgene_12260). pLC-ZsGreen-P2A-Puro and pLC-ZsGreen-P2A-Hygro are available as Addgene plasmids #124302 and #124301 (*17*).

### New Constructs

All primers and gBLOCKs were from IDT, for sequences see Table S1. All inserts and immediate vector context were confirmed by Sanger sequencing (ACGT), except when noted otherwise.

To clone pLC-dCas9VP64-T2A-eGFP, dCas9-VP64 was amplified from lenti dCAS-VP64_Blast (*4*) using primers 4396 and 4397, EGFP was amplified from pLCE (*31*) using primers 4398 and 4395. Products were used for Gibson Assembly with the NheI-EcoRI vector fragment of pLCE.

To clone pXPR_502-sgAAVS1 and pXPR_502-sgCRBN-a1, pXPR_502 was cut using Esp3I (BsmBI, Thermo Fisher Scientific, ER0452) and subjected to T4 DNA ligation with annealed oligos 2692/2693 (AAVS1) or 4684/4685 (CRBN sg-a1, which was picked from the Calabrese library set A).

For Fig.2, we initially constructed an “empty” lentiviral vector pLenti-2xMCS (pL2M) for flexible insertion of promoter-transgene cassettes upstream and downstream of a central polypurine tract (cPPT). For the vector backbone, we cut pLX-sgRNA (*29*) with NotI-HF and NheI-HF to remove a fragment beginning upstream of the Rev response element (RRE) and ending just upstream of the woodchuck hepatitis virus post-transcriptional regulatory element (WPRE). We then re-inserted PCR-amplified fragments containing the RRE (primers 5151 and 5152) and the cPPT (primers 5153 and 5154) using Gibson assembly, resulting in a vector with the following features: RSV/R-U5-Psi-RRE-(*MluI*-*EcoRI*)-cPPT-(*XhoI-NheI*-*AgeI-SalI*) -WPRE-*SalI* -SIN3’LTR. pL2M shares its backbone and LTR sequences with the commonly used sgRNA plasmids pLX-sgRNA (*29*), lentiGuide-Puro (*30*), and pXPR_502 (*11*).

To insert a CMV promoter into the MCS 3’ to the cPPT, L2M was cut using XhoI and NheI-HF. The CMV promoter was amplified from pLCE (*31*) using primers 5157 and 5164. The resulting fragment was digested with XhoI and NheI-HF and ligated into the cut vector using T4 DNA ligase. We named the resulting vector pL2M-CMV. We next inserted a puromycin resistance gene under CMV control in pL2M-CMV to clone pL2M-CMV-Puro. First, pL2M-CMV was cut using NheI-HF and AgeI-HF. The puromycin resistance gene was amplified from lentiGuide-Puro (*30*) using primers 5163/5160, 5183/5184, and 5183/5160. Resulting PCR products were pooled, digested with NheI-HF and AgeI-HF, and ligated into the vector using T4 DNA ligase.

pL2M-CMV-MPH-P2A-Puro and deletion mutants. To insert the MPH-P2A-PuroR fusion proteins under CMV control, pL2M-CMV was cut using NheI-HF and AgeI-HF. A fragment containing a portion of the CMV promoter, and NLS-MCP-linker-SV40-NLS sequences were ordered as gBlock 5169 (IDT). We PCR-amplified a fragment containing codon altered murine p65^AD^ and unaltered human HSF1^AD^ from an unpublished version of lenti MS2-P65-HSF1_Hygro (*4*) that was modified for blasticidin resistance, using primers 5170/5171. We PCR-amplified a fragment containing P2A-puroR from pZIP-P2A-Puro (*18*) using primers 5172/5160. Fragments were joined by Gibson Assembly. In the context of the resulting vector, pL2M-CMV-MPH-P2A-Puro, we deleted the MCP coat protein using primers 5225/5226, the p65^AD^ using primers 5224/5221, the HSF1^AD^ using primers 5220/5249, and p65^AD^-HSF1^AD^ using primers 5220/5221 and the Q5® Site-Directed Mutagenesis Kit (NEB, #E0552S). Resulting mutants were confirmed by full plasmid sequencing (Plasmidsaurus).

To clone pL2M-CMV-MCP-VP64-P2A-Puro, pL2M-CMV-MPH-P2A-Puro was cut using NheI-HF and BamHI-HF. A fragment containing VP64 was PCR amplified from pLC-dCas9VP64-T2A-eGFP using primers 5370/5371. The vector, the PCR amplified fragment, and gBlock 5169 were joined by Gibson Assembly.

To clone pL2M-CMV-MCP-p300^Core^-HA-P2A-Puro, pL2M-CMV-MPH-P2A-Puro was cut using NheI-HF and BamHI-HF and the resulting vector backbone was used for Gibson Assembly with gBlock 5169 (see above) and a fragment containing p300^Core^-HA tag that was PCR amplified from pcDNA-dCas9-p300 (Addgene # 61357) using primers 5372/5373. We introduced the D1399Y mutation using primers 5535/6 and the Q5® Site-Directed Mutagenesis Kit. The resulting mutant was confirmed by full plasmid sequencing (Plasmidsaurus).

To clone pLCM-CMV-MCP-NFZ/NZF-P2A-Puro, we cut pL2M-CMV-MPH-P2A-Puro using NheI-HF and BamHI-HF and used the resulting vector for Gibson Assembly with synthesized fragments 5533 (NFZ) or 5534 (NZF) (Twist Bioscience).

For the sgRNA vectors used in Fig. 4, we excised the PPH-2A-Puro cassette with BamHI and MluI. We used Gibson assembly to insert a puromycin resistance cassette we PCR amplified using primers 5554/5555 and pL2M-CMV-Puro as a template. The resulting vectors are pXPR-Puro-sgAAVS and pXPR-Puro-sgCRBN-a1 (see Fig. S6C for a schematic).

To clone the Tet-ON LV pTO-Zeo-ΔWPRE, we cut pLVX-TetOne-Zeo (*32*) with MluI and NheI and performed Gibson Assembly with a PCR product (primers 5530/2) containing the self-inactivating LTR sequence amplified from pL2M. We next inserted MPH or hrGFP2 cassettes between the EcoRI/AgeI sites of this vector to clone pTO-Zeo-ΔWPRE-MPH or -hrGFP2.

### Production of lentiviral vectors

To produce lentiviral vectors, we co-transfected each transfer vector with pMD2.G and psPAX2 into 293T/17 cells using a 0.624 mg/ml PEI MAX (Polysciences, Catalog # 24765) stock solution (pH 7.4, adjusted using NaOH) and 0.4 pmol DNA/6 well, 3 pmol DNA/10 cm dish, or 7 pmol DNA/15 cm dish, at a ratio of 3.5 μl PEI MAX per 1 μg DNA. A molar ratio of 45% transfer vector, 35% psPax2, and 20% MD2.G was used, except for Fig. 1 and Fig. S1, where all toxic vectors were packaged using 15% transfer vector, ∼54% psPax2, ∼31% pMD2.G, since we found that reducing the amount of transfer vector increases pXPR_502-based lentivirus titers (compare RNA-based titers in Fig. 1D, where 15% transfer vector was used, to those in Fig.2D, where 45% transfer vector was used, p=0.04, ∼1.8x improved titer with 15%). This observation is likely explained by reduced vector toxicity when a lower amount is transfected. Culture media were changed ∼4 hours after transfection. Approximately 72 hours after transfection, we filtered supernatants through 450nm pore size filters and froze aliquots at -80°C. For Fig. 1, lentivirus was first concentrated by ultracentrifugation (Beckman SW 32 Ti rotor, 80,000 x g, 1 hour, 4°C), pellets were incubated with Opti-MEM (Gibco, 31985070) on a platform shaker for at least one hour at 4°C, resuspended by pipetting up and down 20-25 times, and then frozen in aliquots.

### Lentiviral titration

For lentivirus titration by qRT-PCR, we used the LentiX qRT-PCR kit (Takara) according to the manufacturer’s instructions. For functional titration by flow cytometry (FACS) or cell counting, serial dilutions of lentiviral stocks were used to transduce target cells in the presence of 5 μg/ml polybrene. For A375, we plated 15,500 cells/cm^2^ the afternoon before transduction. For BC-3 or BC-3 dCas9-VP64, we split cultures to 3-5 × 10^5^ cells/ml the day before transduction, to ensure robustly proliferating cultures. The next day, BC-3 or BC-3 dCas9-VP64 were adjusted to 3 × 10^5^ cells/ml and ∼0.208 ml/cm^2^. For GFP-based titration, FACS was performed on a BD FACS Canto II, two days after titration with (A375) or without (BC-3) changing media the day after transduction. For functional titration in A375, we changed the culture medium ∼24 hours after transduction to medium containing 1 mg/ml puromycin, maintaining unselected and selected untransduced controls. For functional titration in BC-3, we added 1 μg/ml puromycin without changing the medium, maintaining unselected and selected untransduced controls. 24-30 hours later, when no viable cells remained in the selected untransduced control well, we counted the surviving cells using trypan blue exclusion assay and flow cytometry (ZsGreen controls) and calculated the percentage of live or GFP-positive relative to the untransduced and unselected control. Functional titers were calculated from 4-20% of surviving or GFP-positive cells, assuming a single transduction event per cell.

### Lentiviral transductions

For all transductions, cell numbers and media volumes were scaled approximately by surface area. LVs were added at the indicated MOIs. 24 hours after transduction, A375 were split 1:2 and at the same time selected using 1 μg/ml puromycin. Selecting A375 on day 2 after transduction independently of splitting did not result in improved survival of activator-domain transduced cells (not shown). BC-3 or BC-3/dCas9-VP64 were collected by low-speed centrifugation and resuspended in new medium containing 1 μg/ml puromycin or 300 μg/ml hygromycin. Cell Titer Glo 2.0 (Promega) was used as instructed at the time points indicated in the manuscript, upon completion of selection, determined using a selected untransduced control sample.

### Establishment of BC-3-dCas9-VP64

pLC-dCas9VP64-T2A-eGFP LV was produced as described above and used to transduce BC-3 cells at ∼MOI 0.6, resulting in ∼45% GFP positive cells. We sorted the top 20% GFP expressors using a FACS Aria system, obtaining 4.4×10^5^ live cells that were grown out into the cell pool that was used here (BC-3/dCas9-VP64).

### Growth Curve Analyses in BC-3

BC-3 or BC-3/dCas9-VP64 were transduced and selected as outline above. The unselected and untransduced control cells were typically counted and split the next day, while other samples were first analyzed and passaged 3 (puromycin) or 4 (hygromycin) days after transduction, when selection was complete. At the first passage after selection, all samples were centrifuged, and cells were resuspended in medium without puromycin or hygromycin. From this time point onwards, untransduced and unselected control cells were split together with transduced cell pools. Growth curve analysis was done using Cell Titer Glo 2.0 and resulting values were normalized to cell counts obtained by trypan blue exclusion assay and manual counting of control samples at each passage. At each passage, all samples were adjusted to 3×10^5^ cells/ml, by either diluting or concentrating samples. For cumulative growth curve analyses in Fig. 1G and Fig. S1A, cell counts at each passage were multiplied by all previous dilution factors, prior to normalization to numbers from the control cell pool.

### Western Blots

Cells were washed with cold PBS and lysed with RIPA containing protease inhibitor cocktail (*16*). For A375, cells were scraped into RIPA buffer without prior detachment. For BC-3, cells were collected by low-speed centrifugation. For D3 lysates, cultures were subjected to a dead-cell removal step using the Miltenyi Biotec Dead Cell Removal Kit (Order no. 130-090-101), MS columns, and an OctoMACS separator, all as instructed, since cultures contained large numbers of dead cells right after low MOI transduction and selection. Day 12 lysates did not contain a substantial number of dead cells and were collected directly. After 15 min lysis on ice, lysates were sonicated for 6 cycles (30 seconds on, 30-40 seconds off), cleared by centrifugation, and quantified using BCA assay. Equal amounts of total protein were separated on 4-12% Bis-Tris gels and transferred to nitrocellulose membranes. Membranes were blocked for one hour or overnight in TBS containing 5% non-fat milk powder. Primary antibodies were used at 1:1000 dilutions in TBS containing 0.1% Tween (TBST) and 5% non-fat milk powder. Membranes were washed 3x 15 minutes in TBST, incubated with IRDye-800-conjucated secondary antibodies (LI-COR) at 1:10,000 in TBST containing 5% non-fat milk powder. The following primary antibodies were used: rabbit anti-HSF1 (Cell Signaling Technology 4356), rabbit anti acetylated-Lysine (Cell Signaling Technology 9441), rabbit anti-Enterobacterio Phage MS2 Coat Protein (Millipore-Sigma, ABE76-I), rabbit anti-CRBN (sigma-Aldrich, HPA045910, Fig. 1I), rabbit anti-CRBN (clone F4I7F, Cell Signaling Technology 60312, Fig. 4D), mouse anti-α-tubulin (Cell Signaling Technology 3873). Western Blots were imaged on LI-COR Odyssey FC or LI-COR M Imagers.

### Statistical Analysis

Statistical analyses were done in Graphpad Prism 10, using two-tailed unpaired t tests, and considering p<0.05 as significant, unless indicated otherwise for specific analyses. Numbers of independent repeats are indicated for each experiment.

## Acknowledgments

Flow Cytometry Cell Sorting was performed on a BD FACSAria SORP system, purchased through the support of NIH 1S10OD011996-01 and 1S10OD026814-01. AM, SJ, and ZL were enrolled in the Northwestern University Master of Biotechnology program for part of this study. We thank Dr. Marc Mendillo for the initial gift of anti-HSF1, Drs. Michael C. Bassik and Lacramioara Bintu for sharing the unpublished NFZ/NZF sequences and Drs. Bintu, Mendillo, and Mazhar Adli for helpful discussions and feedback on this manuscript.

## Author contributions

Conceptualization: AM, SJ, EG

Methodology: AM, ZL, SJ, JT, AP, MM, EG

Investigation (data and reagents shown): AM, ZL, SJ, AP, MM, EG

Investigation (unpublished, but informing the study): AP, MM, SR, EG

Visualization: AM, EG

Supervision: EG

Writing—original draft: EG

Writing—review & editing: all authors

## Data and materials availability

All data are available in the main text or the supplementary materials. Any new vectors are available on reasonable request to EG.

## References

1. L. Bendixen, T. I. Jensen, R. O. Bak, CRISPR-Cas-mediated transcriptional modulation: The therapeutic promises of CRISPRa and CRISPRi. Mol Ther 31, 1920–1937 (2023).

2. L. S. Qi, M. H. Larson, L. A. Gilbert, J. A. Doudna, J. S. Weissman, A. P. Arkin, W. A. Lim, Repurposing CRISPR as an RNA-guided platform for sequence-specific control of gene expression. Cell 152, 1173–1183 (2013).

3. L. A. Gilbert, M. H. Larson, L. Morsut, Z. Liu, G. A. Brar, S. E. Torres, N. Stern-Ginossar, O. Brandman, E. H. Whitehead, J. A. Doudna, W. A. Lim, J. S. Weissman, L. S. Qi, CRISPR-mediated modular RNA-guided regulation of transcription in eukaryotes. Cell 154, 442–451 (2013).

4. S. Konermann, M. D. Brigham, A. E. Trevino, J. Joung, O. O. Abudayyeh, C. Barcena, P. D. Hsu, N. Habib, J. S. Gootenberg, H. Nishimasu, O. Nureki, F. Zhang, Genome-scale transcriptional activation by an engineered CRISPR-Cas9 complex. Nature 517, 583–588 (2015).

5. A. W. Cheng, H. Wang, H. Yang, L. Shi, Y. Katz, T. W. Theunissen, S. Rangarajan, C. S. Shivalila, D. B. Dadon, R. Jaenisch, Multiplexed activation of endogenous genes by CRISPR-on, an RNA-guided transcriptional activator system. Cell Res 23, 1163–1171 (2013).

6. L. A. Gilbert, M. A. Horlbeck, B. Adamson, J. E. Villalta, Y. Chen, E. H. Whitehead, C. Guimaraes, B. Panning, H. L. Ploegh, M. C. Bassik, L. S. Qi, M. Kampmann, J. S. Weissman, Genome-Scale CRISPR-Mediated Control of Gene Repression and Activation. Cell 159, 647–661 (2014).

7. P. Mali, J. Aach, P. B. Stranges, K. M. Esvelt, M. Moosburner, S. Kosuri, L. Yang, G. M. Church, CAS9 transcriptional activators for target specificity screening and paired nickases for cooperative genome engineering. Nat Biotechnol 31, 833–838 (2013).

8. M. E. Tanenbaum, L. A. Gilbert, L. S. Qi, J. S. Weissman, R. D. Vale, A protein-tagging system for signal amplification in gene expression and fluorescence imaging. Cell 159, 635–646 (2014).

9. A. Chavez, J. Scheiman, S. Vora, B. W. Pruitt, M. Tuttle, P. R. I. E S. Lin, S. Kiani, C. D. Guzman, D. J. Wiegand, D. Ter-Ovanesyan, J. L. Braff, N. Davidsohn, B. E. Housden, N. Perrimon, R. Weiss, J. Aach, J. J. Collins, G. M. Church, Highly efficient Cas9-mediated transcriptional programming. Nat Methods 12, 326–328 (2015).

10. A. Chavez, M. Tuttle, B. W. Pruitt, B. Ewen-Campen, R. Chari, D. Ter-Ovanesyan, S. J. Haque, R. J. Cecchi, E. J. K. Kowal, J. Buchthal, B. E. Housden, N. Perrimon, J. J. Collins, G. Church, Comparison of Cas9 activators in multiple species. Nat Methods 13, 563–567 (2016).

11. K. R. Sanson, R. E. Hanna, M. Hegde, K. F. Donovan, C. Strand, M. E. Sullender, E. W. Vaimberg, A. Goodale, D. E. Root, F. Piccioni, J. G. Doench, Optimized libraries for CRISPR-Cas9 genetic screens with multiple modalities. Nat Commun 9, 5416 (2018).

12. L. Holtzman, C. A. Gersbach, Editing the Epigenome: Reshaping the Genomic Landscape. Annu Rev Genomics Hum Genet 19, 43–71 (2018).

13. S. Sajwan, M. Mannervik, Gene activation by dCas9-CBP and the SAM system differ in target preference. Sci Rep 9, 18104 (2019).

14. I. B. Hilton, A. M. D’Ippolito, C. M. Vockley, P. I. Thakore, G. E. Crawford, T. E. Reddy, C. A. Gersbach, Epigenome editing by a CRISPR-Cas9-based acetyltransferase activates genes from promoters and enhancers. Nat Biotechnol 33, 510–517 (2015).

15. N. Kuehnle, S. M. Osborne, Z. Liang, M. Manzano, E. Gottwein, CRISPR screens identify novel regulators of cFLIP dependency and ligand-independent, TRAIL-R1-mediated cell death. Cell Death Differ 30, 1221–1234 (2023).

16. M. Manzano, A. Patil, A. Waldrop, S. S. Dave, A. Behdad, E. Gottwein, Gene essentiality landscape and druggable oncogenic dependencies in herpesviral primary effusion lymphoma. Nat Commun 9, 3263 (2018).

17. A. Patil, M. Manzano, E. Gottwein, Genome-wide CRISPR screens reveal genetic mediators of cereblon modulator toxicity in primary effusion lymphoma. Blood Adv 3, 2105–2117 (2019).

18. A. Patil, M. Manzano, E. Gottwein, CK1alpha and IRF4 are essential and independent effectors of immunomodulatory drugs in primary effusion lymphoma. Blood 132, 577–586 (2018).

19. L. Arvanitakis, E. A. Mesri, R. G. Nador, J. W. Said, A. S. Asch, D. M. Knowles, E. Cesarman, Establishment and characterization of a primary effusion (body cavity-based) lymphoma cell line (BC-3) harboring kaposi’s sarcoma-associated herpesvirus (KSHV/HHV-8) in the absence of Epstein-Barr virus. Blood 88, 2648–2654 (1996).

20. J. Joung, S. Konermann, J. S. Gootenberg, O. O. Abudayyeh, R. J. Platt, M. D. Brigham, N. E. Sanjana, F. Zhang, Genome-scale CRISPR-Cas9 knockout and transcriptional activation screening. Nat Protoc 12, 828–863 (2017).

21. K. Omachi, J. H. Miner, Comparative analysis of dCas9-VP64 variants and multiplexed guide RNAs mediating CRISPR activation. PLoS One 17, e0270008 (2022).

22. L. M. Lasko, C. G. Jakob, R. P. Edalji, W. Qiu, D. Montgomery, E. L. Digiammarino, T. M. Hansen, R. M. Risi, R. Frey, V. Manaves, B. Shaw, M. Algire, P. Hessler, L. T. Lam, T. Uziel, E. Faivre, D. Ferguson, F. G. Buchanan, R. L. Martin, M. Torrent, G. G. Chiang, K. Karukurichi, J. W. Langston, B. T. Weinert, C. Choudhary, P. de Vries, J. H. Van Drie, D. McElligott, E. Kesicki, R. Marmorstein, C. Sun, P. A. Cole, S. H. Rosenberg, M. R. Michaelides, A. Lai, K. D. Bromberg, Discovery of a selective catalytic p300/CBP inhibitor that targets lineage-specific tumours. Nature 550, 128–132 (2017).

23. J. Tycko, M. V. Van Aradhana, N. DelRosso, D. Yao, X. Xu, C. Ludwig, K. Spees, K. Liu, G. T. Hess, M. Gu, A. X. Mukund, P. H. Suzuki, R. A. Kamber, L. S. Qi, L. Bintu, M. C. Bassik, Development of compact transcriptional effectors using high-throughput measurements in diverse contexts. bioRxiv, 2023.2005.2012.540558 (2023).

24. S. F. Schmidt, B. D. Larsen, A. Loft, S. Mandrup, Cofactor squelching: Artifact or fact? Bioessays 38, 618–626 (2016).

25. C. Matis, P. Chomez, J. Picard, R. Rezsohazy, Differential and opposed transcriptional effects of protein fusions containing the VP16 activation domain. FEBS Lett 499, 92–96 (2001).

26. M. A. D. Silveira, S. Bilodeau, Defining the Transcriptional Ecosystem. Mol Cell 72, 920–924 (2018).

27. C. H. Ludwig, A. R. Thurm, D. W. Morgens, K. J. Yang, J. Tycko, M. C. Bassik, B. A. Glaunsinger, L. Bintu, High-throughput discovery and characterization of viral transcriptional effectors in human cells. Cell Syst 14, 482–500 e488 (2023).

28. J. Tycko, N. DelRosso, G. T. Hess Aradhana, A. Banerjee, A. Mukund, M. V. Van, B. K. Ego, D. Yao, K. Spees, P. Suzuki, G. K. Marinov, A. Kundaje, M. C. Bassik, L. Bintu, High-Throughput Discovery and Characterization of Human Transcriptional Effectors. Cell 183, 2020–2035 e2016 (2020).

29. T. Wang, J. J. Wei, D. M. Sabatini, E. S. Lander, Genetic screens in human cells using the CRISPR-Cas9 system. Science 343, 80–84 (2014).

30. N. E. Sanjana, O. Shalem, F. Zhang, Improved vectors and genome-wide libraries for CRISPR screening. Nat Methods 11, 783–784 (2014).

31. J. Zhang, D. D. Jima, C. Jacobs, R. Fischer, E. Gottwein, G. Huang, P. L. Lugar, A. S. Lagoo, D. A. Rizzieri, D. R. Friedman, J. B. Weinberg, P. E. Lipsky, S. S. Dave, Patterns of microRNA expression characterize stages of human B-cell differentiation. Blood 113, 4586–4594 (2009).

32. A. F. Castaneda, B. A. Glaunsinger, The Interaction between ORF18 and ORF30 Is Required for Late Gene Expression in Kaposi’s Sarcoma-Associated Herpesvirus. J Virol 93, (2019).

