## Supplemental Figures for "Cytotoxicity of Activator Expression in CRISPR-based Transcriptional Activation Systems"

**This PDF file includes:**
Figs. S1 to S6

**Other Supplementary Materials for this manuscript include the following:**
Table S1 Containing sequences of oligonucleotide and DNA fragments

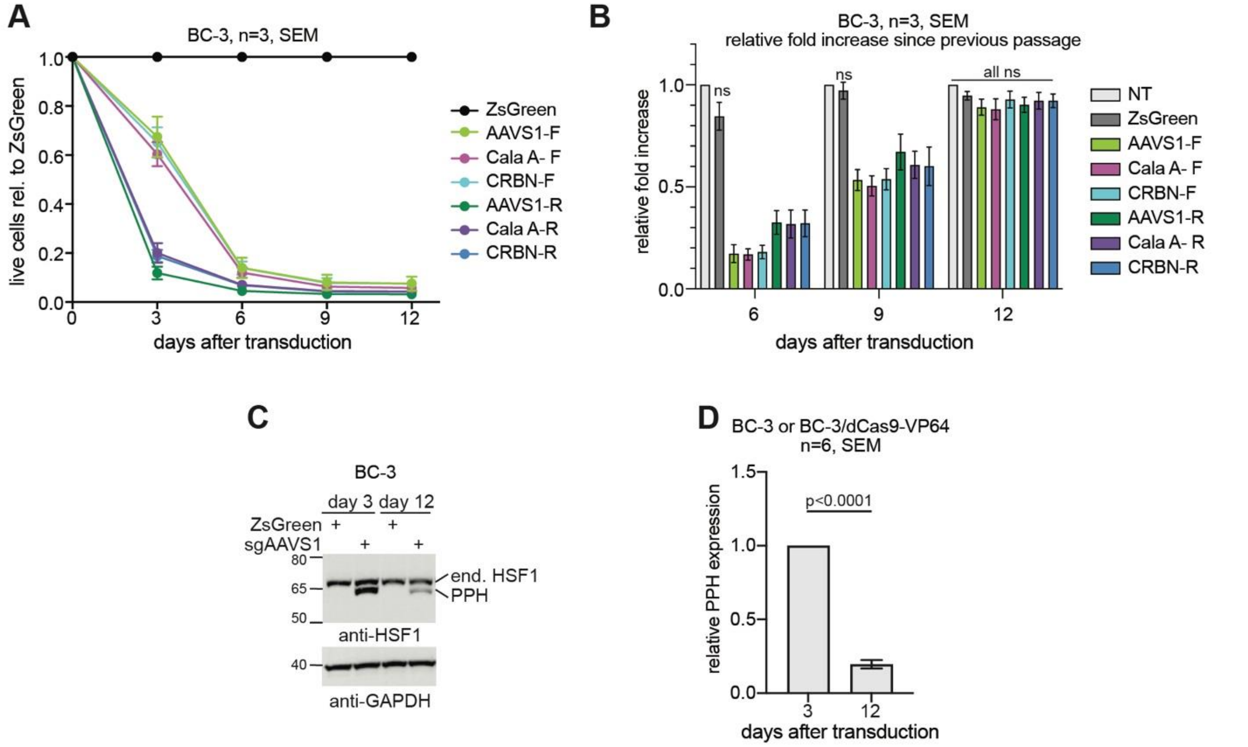


**Fig. S1 Extended Data for Fig. 1.**

**(A)** As in Fig. 1 G, but using parental BC-3 cells. 3 independent repeats, using the 3 LV preps from Fig. 1, panels D-F. All values differed significantly from the ZsGreen control, unpaired t test, $p<0.05$, n=3 independent repeats. Error bars represent SEM.

**(B)** As in Fig. 1 H, but using parental BC-3 cells. Fold increase over the previous passage on day 6, 9, and 12, from the same experiments shown in panel A. Values were normalized to the untransduced and unselected control samples (NT) in this analysis. Values differed significantly from NT at each time point unless specified by ns, unpaired t test, $p<0.05$. Error bars represent SEM.

**(C)** Western Blot analysis of PPH expression as in Fig. 1I but using parental BC-3.

**(D)** Quantification of PPH expression from results as in Fig. 1I and S1C, combining 2 replicates each for pXPR-sgAAVS1 in BC-3, and Calabrese A or pXPR-sgCRBN-a1 in BC-3/dCas9-VP64. Since these vectors behaved similarly in both cell types (Fig. 1G-H, and S1 A-B), we treated the Westerns as six independent replicates for this analysis.

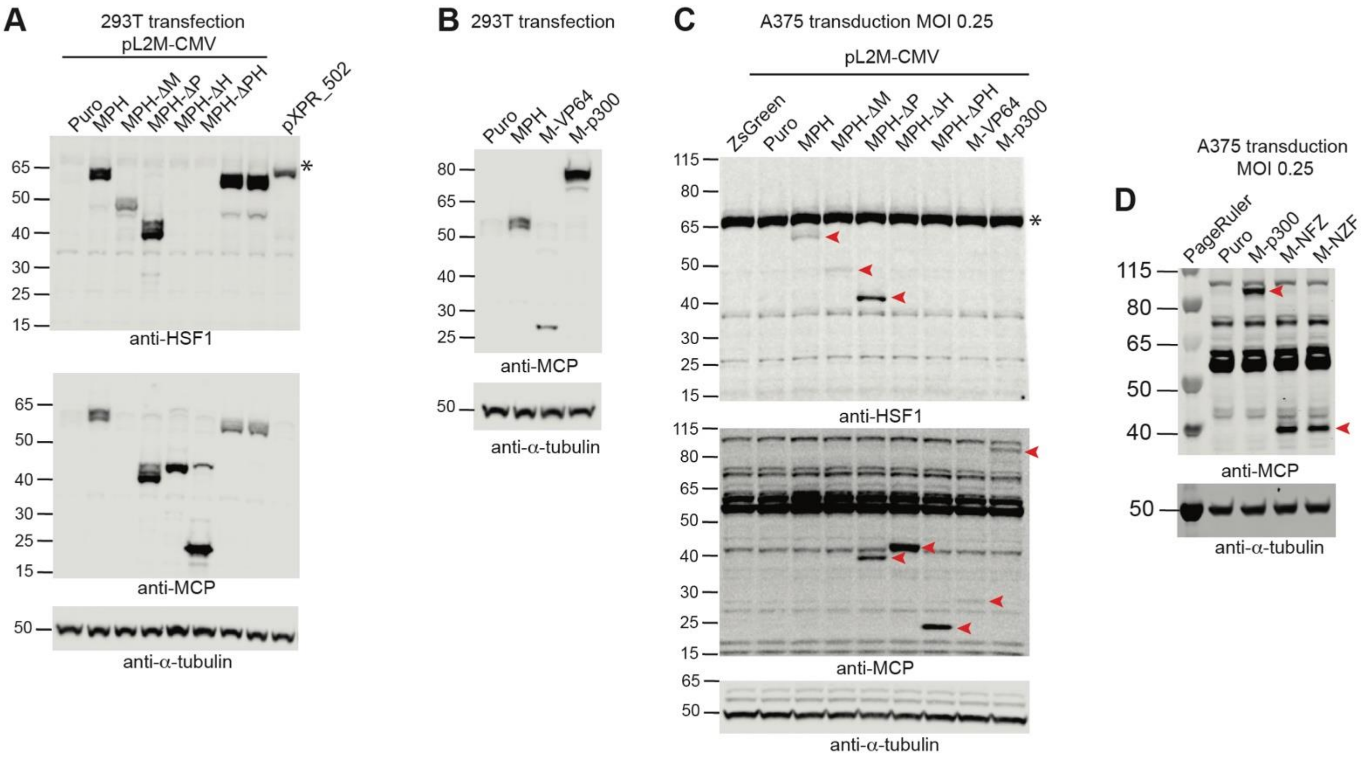


**Fig. S2 Extended Data for Fig. 2.**

**(A)** Western Blot analysis of a subset of the LVs from Fig. 2A-D, after expression in 293T. Lysates were probed for the $HSF1^{AD}$ using anti-HSF1 antibodies, for MCP, and for α-tubulin as a loading control. The likely migration of endogenous HSF1 is marked by an asterisk. Representative of at least 3 independent repeats per vector. Two unlabeled lanes analyze additional vectors that are not described here.

**(B)** Western Blot Analysis of a second subset of the LVs from Fig. 2A-D, after expression in 293T. Lysates were probed for MCP, and α-tubulin served as a loading control. Representative of at least 3 independent repeats per vector.

**(C-D)** As in Fig. S2A-B but using lysates from two days after transduction of A375 at MOI 0.25, without selection. The background is much higher compared to Fig. S2A-B, due to low copy transduction and the background of untransduced cells in the absence of puromycin selection. Toxic protein expression is likely weaker in this experiment due to the beginning loss of transduced cells before harvest. Representative of 2-3 repeats.

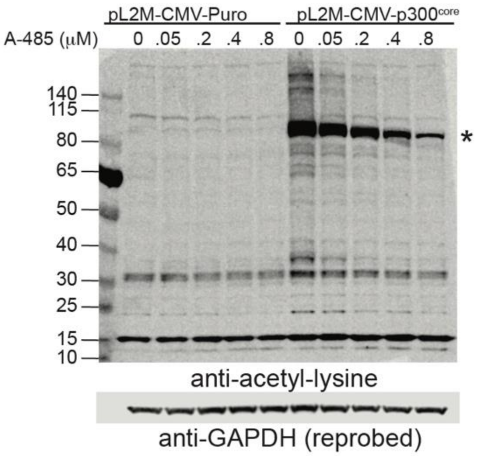


**Fig. S3 Extended Data for Fig. 2E.**
Western Blot analysis of lysine acetylation two days after transduction of A375 at MOI 0.25, without puromycin selection, but otherwise as in Fig. 2E. Reduction of baseline acetylation in untransduced cells over two days was minimal using A-485 concentrations tolerated in A375. We note that off-target acetylation is detected against the background of a majority of untransduced cells in this experiment, due to low MOI transduction. The asterisk marks the location of likely auto-acetylated MCP-$p300^{core}$.

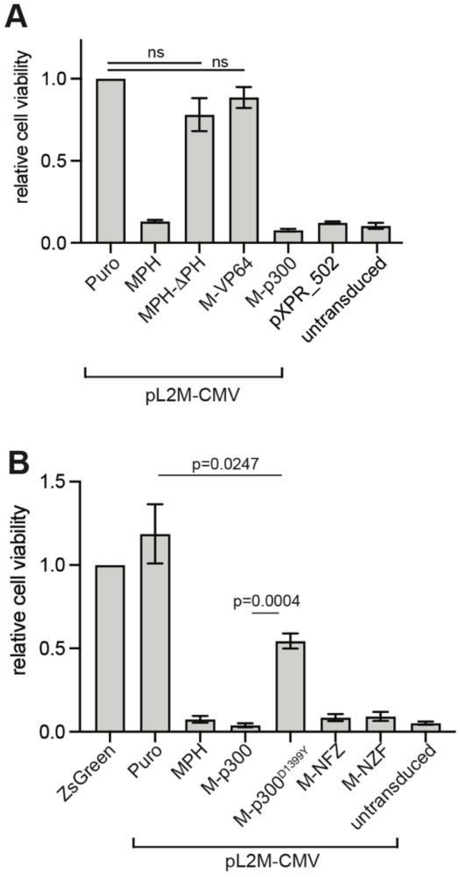


**Fig. S4 Extended Data for Fig. 2.**

**(A)** Like Fig. 2A, but in BC-3 and including only a subset of the vectors. Relative numbers of BC-3 cells surviving puromycin selection after transduction with the indicated vectors at MOI 0.25 based on LV-gRNA content and the functional titer of the ZsGreen control vector, which was not included in the final experiment. Survival was measured using Cell titer Glo 2.0 on day 3 after transduction, and values were normalized to the CMV-Puro control. Values differed significantly from the CMV-Puro control unless indicated by ns, unpaired t test, $p<0.05$, n=3. In this experiment, viability after transductions with lentiviruses expressing MPH, MCP-p300$^{Core}$, and pXPR_502 was not significantly different from untransduced, selected controls (“selected”). **(B)** As in panel A with additional vectors. Values for pL2M-CMV vectors differed significantly from the CMV-Puro control, but, except for MCP-p300$^{Core/D1339Y}$, not from the untransduced, selected control, unpaired t test, $p<0.05$, n=3.

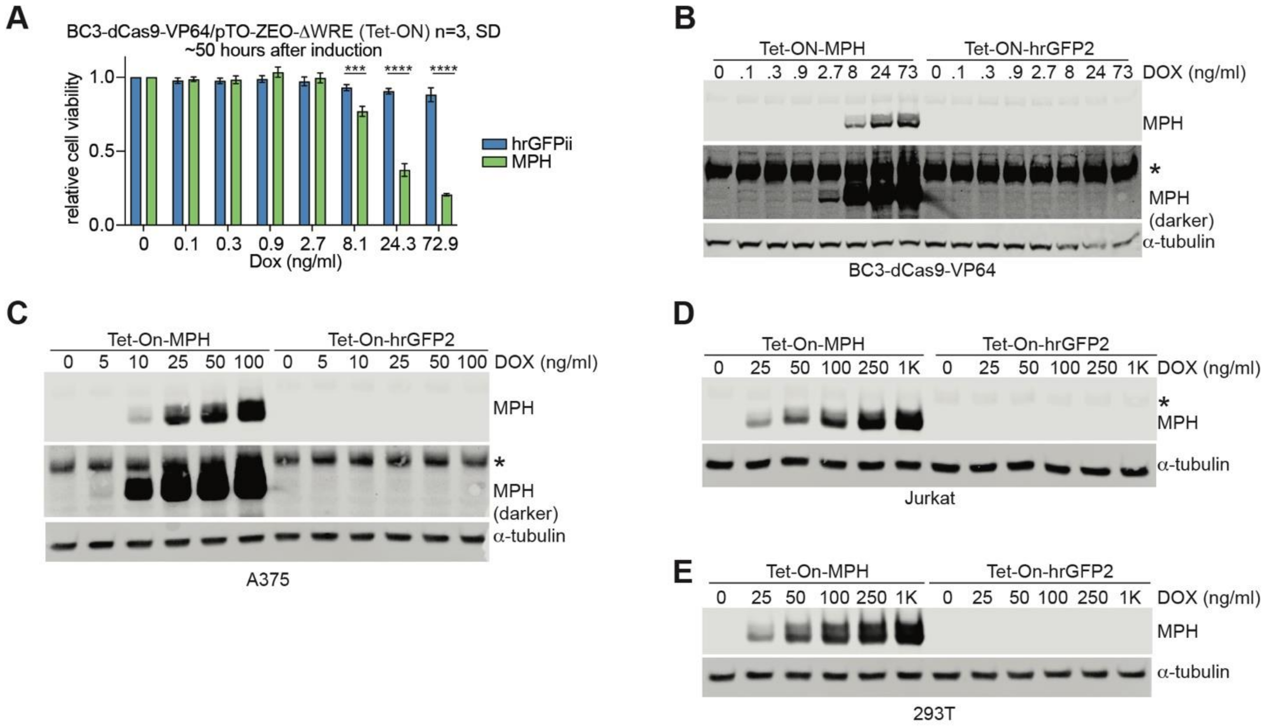


**Fig. S5 Extended Data for Fig. 3.**
**(A)** Experiment as in Fig. 3B, but in BC-3/dCas9-VP64.
**(B)** Western Blot analysis of MPH expression in samples as in Fig. S5A. α-tubulin served as a loading control. n=1, since this panel simply confirms MPH expression. For analysis over replicates see Fig. 4B. The panel shows two different displays of the MPH panel to visualize MPH expression over the range of concentrations.
**(C-E)** As in panel B, but in A375, Jurkat, and 293T cells.

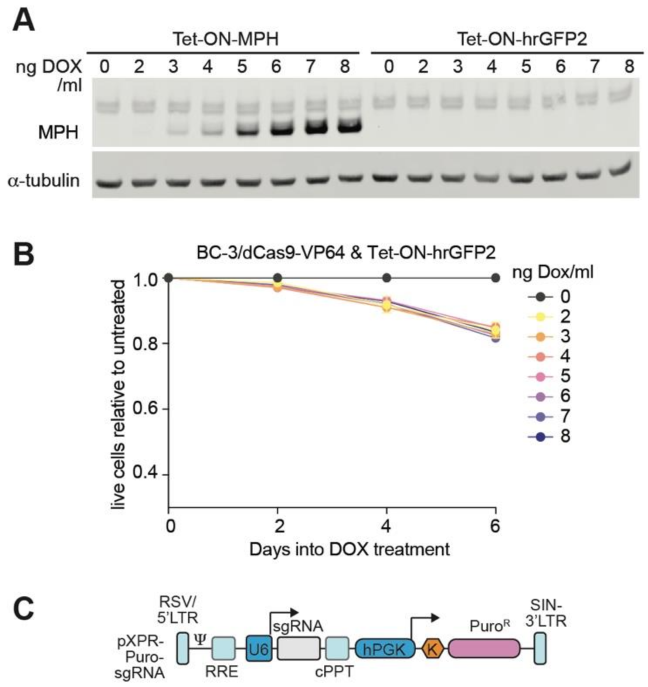


**Fig. S6 Extended Data for Fig. 4**
**(A)** Full panel showing all samples from the experiment in Fig. 4A. Representative of n=3.
**(B)** As in Fig. 4C, but showing results from Tet-ON-hrGFP2.
**(C)** Schematic of the sgRNA vector used in Figs. 4 and S6.
